# A high-throughput, microplate reader-based method to monitor *in vitro* HIV latency reversal in the absence of flow cytometry

**DOI:** 10.1101/2024.09.11.612557

**Authors:** Chantal Emade Nkwelle, Unique Stephens, Kimberly Liang, Joel Cassel, Joseph M. Salvino, Luis J. Montaner, Roland N. Ndip, Seraphine N. Esemu, Fidele Ntie-Kang, Ian Tietjen

**Author notes:** **Correspondence:** Fidele Ntie-Kang [Center for Drug Discovery, Faculty of Science, P.O. Box 63, University of Buea, Buea, Cameroon] and Ian Tietjen [The Wistar Institute, 3601 Spruce Street, Philadelphia, PA 19104, USA] (F.N.K.); (I.T.) (F.N.K.); (I.T.).

## Abstract

J-Lat cells are derivatives of the Jurkat CD4+ T cell line that contain a non-infectious, inducible HIV provirus with a GFP tag. While these cells have substantially advanced our understanding of HIV latency, their use by many laboratories in low and middle-income countries is restricted by limited access to flow cytometry. To overcome this barrier, we describe a modified J-Lat assay using a standard microplate reader that detects HIV-GFP expression following treatment with latency-reversing agents (LRAs). We show that HIV reactivation by control LRAs like prostratin and romidepsin is readily detected with dose dependence and with significant correlation and sensitivity to standard flow cytometry. For example, 10 µM prostratin induced a 20.1 ± 3.3-fold increase in GFP fluorescence in the microplate reader assay, which corresponded to 64.2 ± 5.0% GFP-positive cells detected by flow cytometery. Similarly, 0.3 µM prostratin induced a 1.7 ± 1.2-fold increase compared to 8.7 ± 5.7% GFP-positive cells detected. Using this method, we screen 79 epigenetic modifiers and identify molibresib, quisinostat, and CUDC-101 as novel LRAs. This microplate reader-based method offers accessibility to researchers in resource-limited regions to work with J-Lat cells and more actively participate in global HIV cure research efforts.

**Highlights:**

- J-Lat T-cell lines are important to HIV cure research but require flow cytometry
- We describe a method to work with J-Lat cells using a standard microplate reader
- This assay can detect control LRAs similar to flow cytometry and discover new LRAs
- This assay allows low-resourced laboratories to contribute to HIV cure research

## 1. Introduction

Current antiretroviral therapy (ART) does not eliminate latent HIV proviruses present in CD4+ T-cells that can reactivate at any time to produce infectious virus. As a result, people living with HIV must maintain an ART regimen for life. Successfully identifying and eliminating these latent HIV reservoirs within CD4+ T-cells would mark a major milestone toward achieving ART-free HIV remission or an HIV cure. A common approach in early-stage HIV cure-based drug discovery is to identify small molecules that can reactive HIV expression from latently infected cells as part of larger efforts to eliminate these cells through cell death or immune system induction. This approach, frequently termed “shock-and-kill” or “kick-and-kill,” has identified hundreds of latency-reversing agents (LRAs) that reactivate virus expression in HIV-infected primary cells, animal models of latent HIV infection, and/or in people living with HIV (Board et al., 2022; Mbonye and Karn, 2024). However, as no LRA-based strategy has demonstrated consistent reduction of HIV reservoirs in humans, optimizing existing LRAs as well as discovery of new LRAs remain critical.

Discovery and validation of new LRAs frequently begin with the use of HIV reservoir-based cell lines which afford high-throughput and low-cost screening of chemical libraries. Among the available cell lines are J-Lat cells, which are derived from Jurkat CD4+ T-cells but contain a latent HIV provirus where the viral *env* and *nef* genes are replaced with a GFP reporter (Jordan et al., 2003). As a result, cells produce GFP upon HIV latency reversal and virus expression, which is readily monitored by flow cytometry. Using this approach, numerous LRAs and other regulators of virus transcription have been discovered, significantly advancing our understandings of HIV latency as well as development of therapeutic leads to eventually achieve ART-free HIV remission or cure (e.g., Abdel-Mohsen et al., 2016; Abner et al., 2018; Hashemi et al., 2018; Richard et al., 2018; Acchioni et al., 2019; Matsuda et al., 2019; Hseih et al., 2023; Nichols Doyle et al., 2023).

One attractive characteristic of J-Lat cell lines, in addition to their open availability, is that they can produce largely intact but non-infectious provirus, which makes them suitable for use by both early-stage trainees and laboratories which lack sufficient biosafety containment for culturing infectious HIV. This may be particularly appealing to many low-resource laboratories in low and middle-income countries like those in Sub-Saharan Africa, which continues to be disproportionately affected by the HIV pandemic (van Schalkwyk et al., 2024) and where supporting more locally based HIV cure research is a high priority. However, while J-Lat cells can be readily cultured in a laboratory with a biosafety cabinet and cell culture incubator, detection of latency reversal in these cells remains challenging for many low-resource labs as they frequently lack access to costly flow cytometry equipment and the ongoing expertise needed to use and maintain it. As a result, variations of standard J-Lat-based assays are needed to make this important research tool more available to those in resource-limited laboratories.

Toward addressing this barrier, we describe here a modified J-Lat based assay which uses a standard microplate reader to detect latency reversal from both J-Lat 10.6 cells, which readily respond to control LRAs by high GFP expression in most cells, as well as J-Lat 6.3 cells, which induce low-level GFP following latency reversal. We show here that comparable and reproducible dose-dependent latency reversal due to control LRAs is detected by microplate reader with similar sensitivity to flow cytometry methods. We also demonstrate proof-of-concept of novel LRA discovery by screening a library of 79 chemical epigenetic modifiers to identify three compounds not previously reported to function as LRAs. This alternative assay approach, which can substitute for flow cytometry, offers improved accessibility to researchers in resource-limited regions to discover and characterize novel LRAs and HIV latency as well as more actively participate in global HIV cure efforts.

## 2. Materials and Methods

### 2.1. Cells and reagents

J-Lat T cell clones 10.6 and 6.3 were obtained from the NIH AIDS Reagent Program, Division of AIDS, NIAID, NIH (contributed by Dr. Eric Verdin; Jordan et al., 2003). Cells were cultured in R10+ medium [RPMI 1640 with HEPES and L-glutamine (Corning Life Sciences, Corning, NY, USA), 10% heat-inactivated fetal bovine serum (Gibco, Thermo Fisher Scientific, Waltham, MA, USA), and 100 units/mL of penicillin plus 100 µg/mL streptomycin (Gibco)] in a humidified incubator at 37 °C and 5% CO_2_. Cells were cultured to a cell density of 1 – 2 million cells/mL before use and subcultured to a concentration of 10^5^ cells/mL every 2 – 3 days. Excess cells were stored in cryopreservation vials (Cryotube vials, Thermo Fisher Scientific) at a concentration of 2 * 10^6^ cells/mL in R10+ supplemented with 40% FBS and 10% DMSO, slow-frozen in a Mr. Frosty cryofreezing container containing isopropanol (Thermo Fisher Scientific) for 24 hours at -80 °C, and stored long-term at - 120 °C or lower. Cultured cells were maintained for a maximum of 30 passages, after which a frozen vial of cells was rapid-thawed and cultured using standard cell culture thawing techniques.

Control LRAs prostratin and romidepsin were purchased from Sigma-Aldrich (St. Louis, MO, USA) and MedChemExpress (Monmouth Junction, NJ, USA), respectively. Epigenetic inhibitors were obtained from the SelleckChem Anti-Cancer Library (Selleck Chemicals, Houston, TX, USA). Control LRAs and test agents were diluted in DMSO and stored at -20 °C until use.

### 2.2. LRA screening

Test agents, control LRAs, and vehicle controls were diluted in PBS (Corning Life Sciences, Corning, NY, USA) and aliquoted in duplicate into single wells of sterile V-bottom or U-bottom 96-well plates (Corning) in 2 µL volumes each reflecting 100X final concentrations. Cell concentrations for J-Lat cell culture were determined using a standard hemocytometer. Cultures were then centrifuged at 500 g for 5 minutes, aspirated of culture medium, and resuspended in fresh, prewarmed R10+ to a concentration of 5 x 10^6^ cells/mL unless otherwise specified. 200 µL aliquots of resuspended cells were then distributed into wells. If cell viability using resazurin was planned, an additional two wells containing only R10+ were also prepared on each plate. Cells were then placed in a humidified incubator at 37 °C and 5% CO_2_ for 24 ± 4 hours.

If flow cytometry was planned, cells were gently resuspended following incubation, and 20 µL of each cell culture was transferred to a fresh 96-well plate. Cells were then supplemented with 180 µL of pre-warmed R10+ medium and optionally incubated for up to 4 additional hours before analysis by flow cytometry. Flow cytometry was performed using a FACSCelesta Flow Cytometer (BD Bioscience, Franklin Lakes, NJ, USA), where gating of live cells and percent HIV latency for each well were obtained as described previously (Richard et al., 2018). Flow cytometry data were analyzed using FlowJo v.10.10.0 software (FlowJo LLC, Ashland, OR, USA).

For microplate reading, following incubation, cell culture plates were spun at 500 g for 5 minutes, flicked over a waste receptable in a smooth motion to discard supernatant, lightly pat-dried against clean paper towels, and resuspended in 200 µL of pre-warmed PBS. Cells were washed an additional three times in pre-warmed PBS using these steps. After the final wash, cells were resuspended in 200 µL PBS and transferred to a white/opaque 96-well flat bottom plate (Thermo Fisher Scientific). GFP fluorescence was then measured using a ClarioStar plate reader (BMG Labtech, Cary, NC, USA).

If measurement of cell viability was planned, 20 µL of a 0.2 mg/mL stock of resazurin (Sigma-Aldrich) was added to each well, and cells were incubated in the white/opaque plate (in PBS) with a lid at 37 °C and 5% CO_2_ for an additional 2 to 4 hours. Cells are then re-read in the microplate reader for absorbance at 570 nm or fluorescence set as close as possible to 550 nm excitation and 590 nm emission. Background signal was then subtracted from all wells based on the signal from control wells containing R10+ medium and resazurin but no cells.

## 3. Results and Discussion

### 3.1. Overview of microplate reader-based LRA screening method

Figure 1 presents an overview of the microplate reader-based method to measure HIV latency reversal from J-Lat cells *in vitro*. To begin, test agents, LRA controls, and vehicle controls (e.g., DMSO if used to dissolve and store LRAs long-term) are diluted in PBS to working concentrations that are 100-fold more concentrated than final concentrations. 2 µL of each test agent or control is then aliquoted in duplicate into single wells of 96-well V-bottom cell culture plates, with the expectation that subsequent addition of J-Lat cells will dilute the concentrations of test agents and controls by 100-fold. This approach is recommended for LRA stocks that are dissolved in DMSO, as we find that DMSO test concentrations greater than 0.6% can affect background GFP expression and/or J-Lat T-cell viability. Each 96-well plate also contains an untreated cell control, a vehicle-only control (e.g., 0.1% DMSO final concentration, or 2 µL of a 10% DMSO solution), and a positive control like prostratin (e.g., 10 µM final concentration, or 2 µL of a 1 mM stock concentration in 10% DMSO) or Phorbol 12-myristate 13-acetate (e.g., 10 nM final concentration, or 2 µL of a 1 µM stock concentration in 10% DMSO). If cell viability measures were desired, an additional two cells containing only medium were added to each plate. Control conditions are also performed in duplicate. We also use V-bottom or U-bottom plates for concentrating cell pellets in centrifuge steps following incubation below.

**Figure 1.**
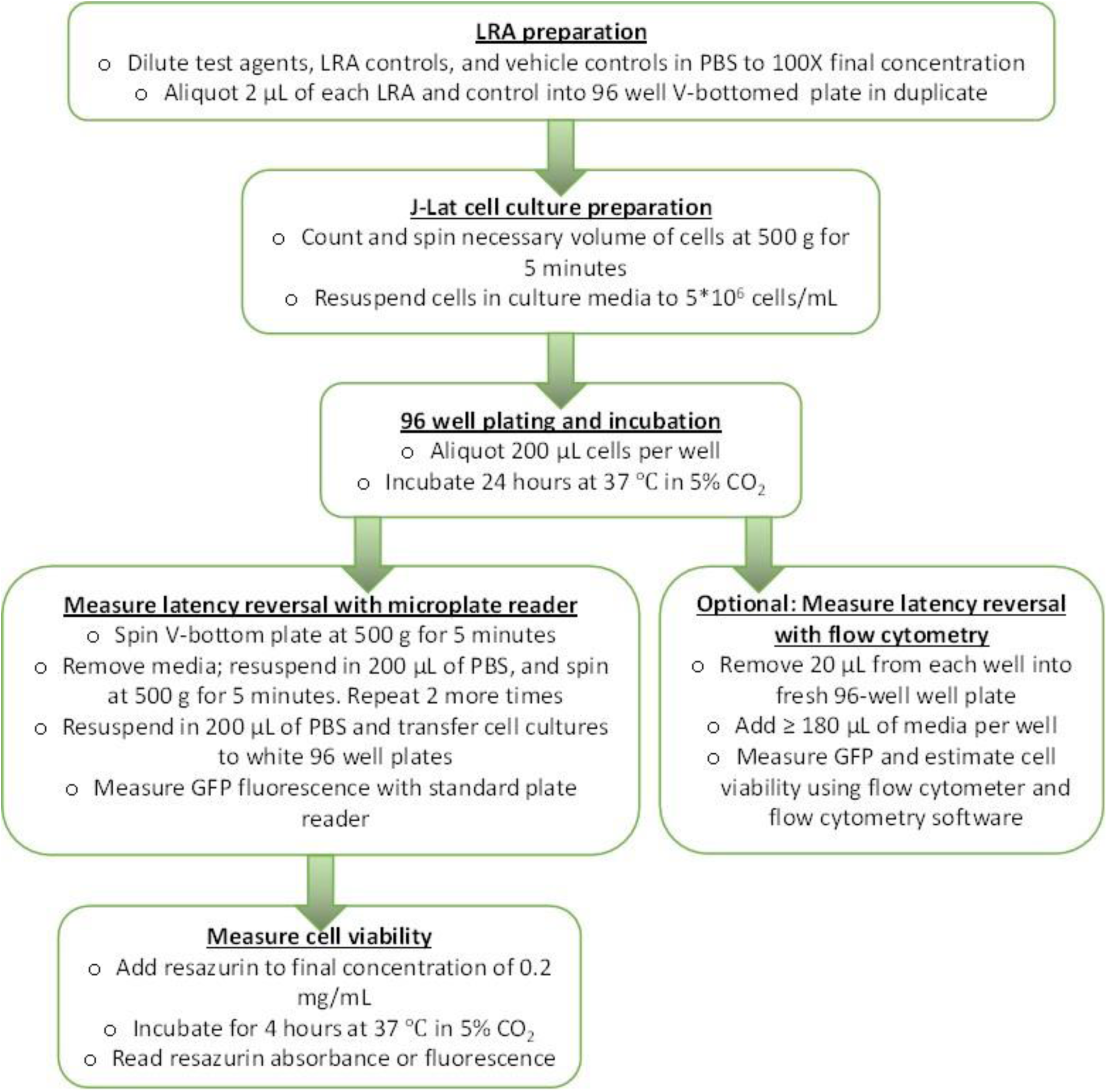
Flowchart of microplate reader-based method using J-Lat cells.

After test agents are aliquoted, J-Lat cell cultures are removed from the incubator and counted using a hemocytometer, and the number of cells required for an experiment is calculated. In our experiments, each well receives 1 million J-Lat cells in a volume of 200 µL (i.e., cells are resuspended in fresh media to a concentration of 5 million cells/mL). For example, if all wells of a 96 well plate are used, a total of 96 million cells resuspended in 19.2 mL of fresh media is needed. In practice, for each 96-well plate, we factor an additional 10% of cells to account for pipetting error, such that each 96-well plate will require 106 million cells, enough for 106 wells, resuspended in 21.2 mL of fresh media. We aliquot 200 µL of cells per well as this dilutes test agents and controls by ∼100-fold to 1X final concentrations. 200 µL cell culture also have minimal evaporation following overnight incubation, thereby reducing evaporation’s effects on cell viability, LRA and control concentrations, and assay reproducibility. As described below, we also found that cells resuspended to a final concentration of 1 million cells/well in a volume of 200 µL, or 5 million cells/mL, best approximated flow cytometry results obtained in parallel. Cell cultures of 100 µL or less can be used to conserve cells and LRAs/controls; however, we then recommend reserving the outer wells of the 96 well plate to fill with > 200 µL PBS to minimize cell culture evaporation. Once these conditions are calculated, the required number of cells are then removed from the parental cell culture, centrifuged at 500 g for 5 minutes at room temperature, and resuspended in fresh media to the desired final concentration, for example 5 million cells/mL. After 200 µL cell aliquots are added to each well (i.e., 1 million cells per well), plated cells are covered with a lid and incubated for 24 hours (± 4 hours) in a humidified incubator at 37 °C with 5% CO_2_.

Following incubation, cell culture plates are centrifuged at 500 g for 5 minutes at room temperature. Cell culture media is removed by inverting the cell culture plate, performing one rapid “flick” of the plate to dispense media into a waste receptacle, and briefly and lightly dabbing the plate on a dry paper towel to remove remaining media. After “flicking,” cell pellets remain adhered to the V-bottom or U-bottom plate along with a few negligible microliters of residual media. Cells are then resuspended in 200 µL of PBS and centrifuged again at 500 g for 5 minutes, after which PBS is removed by flicking; this process is performed a total of three times. Following three washes with PBS, cells are resuspended once more in 200 µL PBS, and resuspended cells are transferred to white 96-well plates. Cells are then immediately read in a 96-well microplate reader that can detect GFP fluorescence. Data are then presented as the fold-change of GFP fluorescence relative to the average GFP fluorescence of untreated or vehicle-treated cells (e.g., 0.1% DMSO). If necessary, cells can be fixed in a final concentration of 4% formalin or paraformaldehyde, sealed with parafilm, and stored at 4 °C in the dark for up to two weeks before measuring GFP; however, this method would obviously preclude downstream measurements of cell viability.

After reading GFP, cells can be optionally measured for cell viability. We usually treat cells with a final concentration of 2 µg/mL resazurin (alamar blue) prepared from a 0.2 mg/mL stock in PBS followed by further incubation in a humidified incubator at 37 °C with 5% CO_2_ for 4 hours. Cells are then re-read in the microplate reader for absorbance at 570 nm or fluorescence set as close as possible to 550 nm excitation and 590 nm emission. While we find that resazurin gives the best reproducibility and sensitivity for measuring cell viability, other reagents like 3-(4,5-dimethylthiazol-2-yl)-2,5-diphenyltetrazolium bromide tetrazolium (MTT) and a standard MTT reduction assay can also be used. For data analysis, the background signal of medium-only wells is then subtracted from all wells, and resulting data are then normalized to the average intensities of untreated or vehicle-treated cells.

### 3.2. Robust and quantitative GFP fluorescence is J-Lat 10.6 cells is detected by microplate reader-based method and correlates with fluorescence detected by flow cytometry

Using this approach, we assessed the ability of the microplate reader-based method to detect HIV reactivation induced by control LRAs in J-Lat 10.6 cells. These LRAs included prostratin, which reverses HIV latency through activating protein kinase C signaling, and romidepsin, which acts via inhibition of histone deacetylases (HDACs) (Andersen et al., 2018). To measure dependence of cell concentration on GFP signals obtained by microplate reader, we treated cells at three different concentrations including 1, 2.5, and 5 * 10^6^ cells/mL (corresponding respectively to 2*10^5^, 5*10^5^, and 10^6^ cells/well in 200 µL final volume) with LRAs. From this experiment, we observed robust GFP detection at all cell concentrations for both LRAs with dose dependence (Figures 2A-C). We also observed that higher cell concentrations induced more detectable GFP signal from both LRAs. For example, we observed a 12.3 ± 1.4-fold increase in GFP fluorescence (mean ± s.e.m.) in cells treated with 10 µM prostratin, relative to cells treated with 0.1% DMSO vehicle control, when cultured at 10^6^ cells/mL (Figure 2A), while cells cultured at 2.5 and 5 * 10^6^ cells/mL respectively exhibited 21.3 ± 2.6 and 20.1 ± 3.3-fold increases in GFP, relative to 0.1% DMSO-treated cells (Figures 2B-C). In contrast, in cells cultured at 10^6^ cells/mL, we observed only a 5.8 ± 0.5-fold increase in GFP fluorescence from cells treated with 0.3 µM romidepsin, although this concentration of romidepsin did induce more GFP fluorescence in cells cultured at 2.5 and 5 * 10^6^ cells/mL (respectively 7.1 ± 0.7 and 9.3 ± 1.5-fold increases in fluorescence, respectively; Figures 2A-C). However, when these cells were subsequently assessed for cell viability following staining with resazurin, we observed, as expected, that while prostratin induced a maximum of 47.2 ± 6.1% loss in viability at 30 µM, substantial cell death was observed in romidepsin-treated cells, with calculated half-maximal cytotoxic concentrations (CC_50_s) of 0.037 µM or lower (Figures 2D-F). This indicates that the reduced romidepsin-dependent signal observed relative to prostratin-dependent fluorescence is attributable to cytotoxicity.

**Figure 2.**
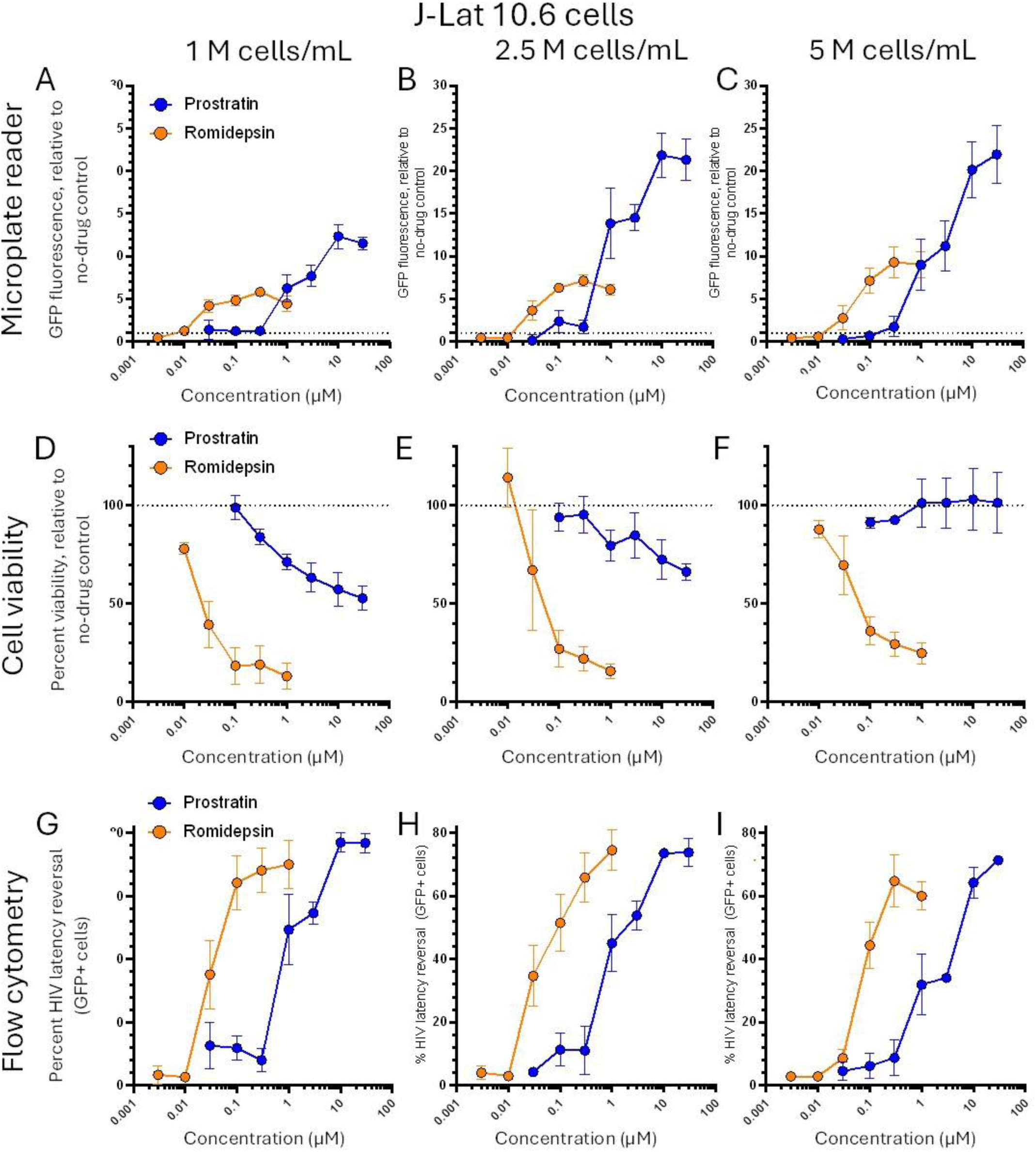
Activity of prostratin and romidepsin in J-Lat 10.6 cells as measured by the microplate reader-based method (**A-C**). Panels **G-I** show viability of the same cells from the microplate reader-based method, while panels **D-F** show GFP expression in these cells as detected by flow cytometery. In columns, **A**, **D**, and **G** are results for J-Lat cells cultured for 24 hours at 1 million cells/mL, **B**, **E**, **H** show cultures at 2.5 million cells/mL, and **C**, **F**, **I** show cultures at 5.0 million cells/mL. All data are presented as the mean ± s.e.m. from four independent experiments.

To compare this microplate reader-based assay with standard methods, we removed 20 µL of each cell culture following the 24-hour incubation above and before plate spinning and PBS washes. These cells were then added to 180 µL of fresh media and processed directly by flow cytometry (Figure 1). Removal of these cells did not obviously affect the magnitude of GFP detection by microplate (data not shown). As shown in Figures 2G-I, similar levels of GFP fluorescence in live-gated cells were observed across all cell concentrations, indicating no major effects on latency reversal due to cell concentration, although somewhat fewer GFP-positive cells were seen at some concentrations of prostratin in cells cultured at 5 x 10^6^ cells/mL (Figure 2I). For example, at 3 µM prostratin we observed 54.7 ± 3.6 and 53.8 ± 4.7% GFP-positive cells from the live cell gates of cells cultured at 1 and 2.5 x 10^6^ cells/mL, respectively, but only 34.1 ± 1.5% GFP-positive cells in the live cell gates from cells cultured at 5 x 10^6^ cells/mL, indicating minor but consistent dampening of GFP expression induced by control LRAs in cells cultured at high concentrations. In contrast, GFP signals induced by romidepsin were robust across all samples where, for example, at 0.3 µM romidepsin, we observed 68.1 ± 7.0, 65.8 ± 7.8, and 61.4 ± 11.6% GFP-positive cells from the live cell gates of cells cultured at 1 and 2.5, and 5 x 10^6^ cells/mL, respectively (Figures 2G-I). Notably, unlike observations from the microplate reader, latency reversal induced by romidepsin also approximated that of prostratin in flow cytometry studies, which reflects standard flow cytometry data analysis where only the subset of live cells, based on characteristic forward and side scatter parameters, are gated and analyzed. As such, screening of chemical libraries using the microplate reader-based method is likely to identify not only LRAs that can induce GFP fluorescence and latency reversal but also select for those that do not cause overt cytotoxicity.

To determine the extent to which HIV latency reversal signals detected by microplate reader (as measured by fold-change in GFP expression relative to no-drug control) correlated to signals detected by flow cytometry (as measured by percent GFP-positive, live-gated cells), we obtained the total data points of J-Lat 10.6 cells treated with prostratin and romidepsin at all concentrations across four independent experiments and graphed microplate reader data as a function of flow cytometry data (Figure 3). By performing simple linear regression analysis, we found that correlations between microplate reader and flow cytometry data improved in proportion to the concentration of J-Lat 10.6 cells in cell culture (Figure 3A-C). For example, while J-Lat 10.6 cells cultured at 1 million cells/mL exhibited an r^2^ value of only 0.32 between the respective microplate reader and flow cytometry data, we observed good correlation between these readings of the same experiments when J-Lat 10.6 cells were cultured at 2.5 million cells/mL (r^2^ = 0.55) and 5 million cells/mL (r^2^ = 0.65). Within the subset of data points where cells were treated with only one control LRA in J-Lat cells cultured at 5 million cells/mL, we also observed good correlation of the two detection methods in cells treated with prostratin (r^2^ = 0.81) and romidepsin (r^2^ = 0.79) (Figure 3D-E). All of these correlations were statistically significant by linear regression (p < 0.001).

**Figure 3.**
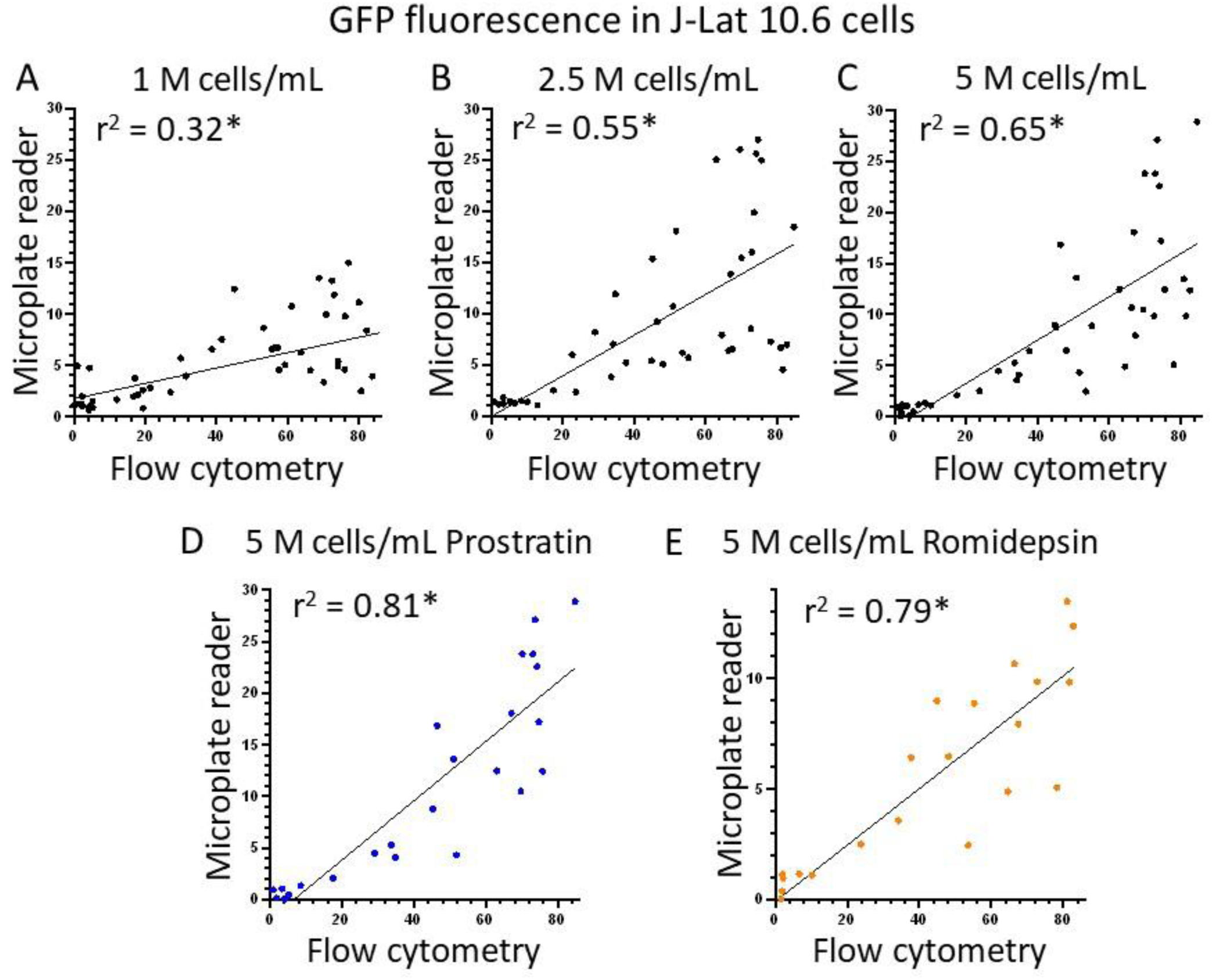
Comparisons of LRA-induced GFP detection using the microplate reader-based method and flow cytometry. Panels **A-C** show correlations of all data points in Figure 2 across four independent experiments for prostratin and romidepsin with microplate reader data graphed on the y-axis (indicating fold-change in GFP expression over untreated cells) and flow cytometry data on the x-axis (indicating percent GFP-positive cells in live cell gate). Panels **A-C** show results from all experiments with 1 million cells/mL, 2.5 million cells/mL, and 5 million cells/mL, respectively. For panel **C**, individual data for prostratin (**D**) and romidepsin (**E**) are shown. *, linear regression p-value < 0.001.

Taken together, these results indicate that the microplate reader-based method detects HIV latency reversal in J-Lat cells by LRAs with dose dependence that recapitulates results obtained by traditional flow cytometry methods. Based on these results, we performed subsequent experiments using J-Lat cells at a concentration of 5 * 10^6^ cells/mL, or 10^6^ cells/well in 200 µL.

### 3.3. Lower but reproducible GFP fluorescence is J-Lat 6.3 cells is detected by microplate reader-based method

To determine whether the microplate reader-based method of detecting HIV latency reversal could be expanded to other J-Lat cell lines, we next investigated whether we could detect GFP expression induced by prostratin or romidepsin in J-Lat 6.3 cells. These cells represent another J-Lat cell clone where the HIV provirus is integrated at a different genomic location and where HIV is poorly induced by LRAs (Fernandez and Zeichner, 2010). However, unlike J-Lat 10.6 cells, where GFP fluorescence of 10 to 20-fold over untreated control cells was routinely detected following romidepsin or prostratin treatment (Figure 2C), we observed no more than a 2.2 ± 0.4-fold increase (mean ± s.e.m.) in fluorescence in the presence of 0.3 µM romidepsin (Figure 4A). Consistent with the low induction of virus expression by LRAs in this cell model, we also detected low levels of GFP-positive cells by flow cytometry, for example following treatment with 10 µM prostratin (average 11.7 ± 5.5% GFP-positive cells) or 0.3 µM romidepsin (average 13.3 ± 6.2% GFP-positive cells; Figure 4B). While dose-response curves for both prostratin and romidepsin were detected in J-Lat 6.3 cells using both methods, more variability between three independent experiments for each assay was observed. Furthermore, when graphing microplate reader and flow cytometry datapoints for two experiments where both assays were performed, a lower correlation was observed between the two assays (r^2^ = 0.37; Figure 4C), although this correlation remained significant (p = 0.005). Finally, while correlations for the subset of datapoints from prostratin treatment had good correlation (r^2^ = 0.67; p = 0.004), no correlation was observed in romidepsin-treated samples (r^2^ = 0.11; p = 0.36), possibly due to the loss of cell viability at high concentrations of romidepsin which in turn may affect low-level GFP expression in J-Lat 6.3 cells (Figure 4D). These results suggest that the microplate reader-based method is capable of detecting HIV latency reversal by LRAs with dose-dependence in J-Lat 6.3 cells but with lower magnitudes and more variability.

**Figure 4.**
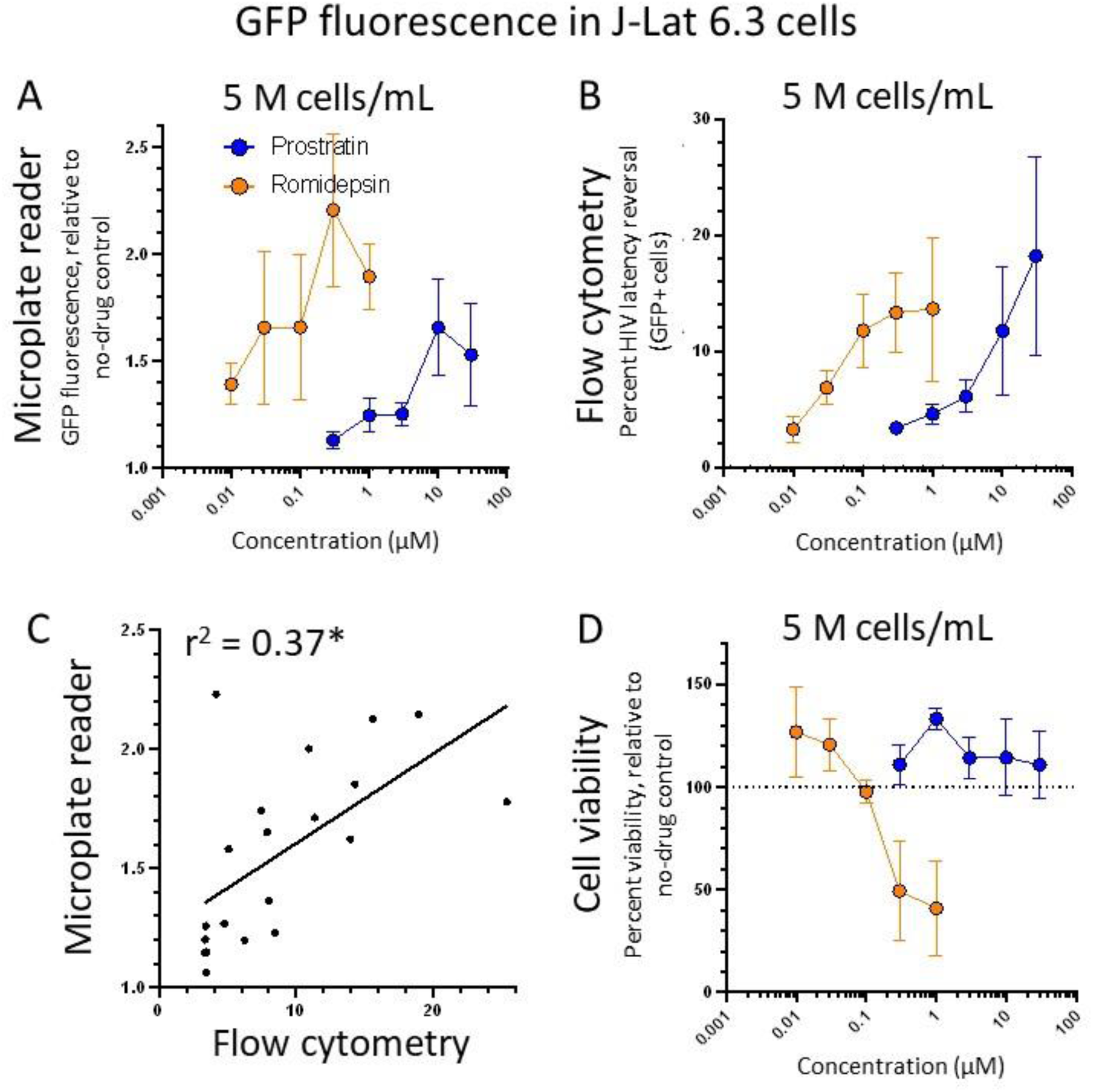
Activity of prostratin and romidepsin in J-Lat 6.3 cells as measured by the microplate reader-based method (**A**) and flow cytometry (**B**). Panel **C** shows correlations of data points with microplate reader data graphed on y-axis (indicating fold-change in GFP expression over untreated cells) and flow cytometry data on x-axis (indicating percent GFP-positive cells in live gate). Panel **D** shows viability of cells from the microplate reader-based method in **A**. All data are presented as the mean ± s.e.m. from three independent experiments except panel **C** which reflects data from two independent experiments. *, linear regression p-value = 0.005.

3.4. *Discovery of novel LRAs in J-Lat 10.6 cells using microplate reader-based method*

To determine whether the microplate reader-based method could be used to identify novel LRAs, we assembled a library of 79 epigenetic modifiers (**Table 1**) and screened them in duplicate at 10 µM in J-Lat 10.6 cells using the procedures above. We also assessed the same cells in parallel by flow cytometry, where good correlation of data from both microplate reader and flow cytometry was maintained (r^2^ = 0.61, p < 0.001; Figure 5A). When focusing exclusively on the microplate reader-based data, we found that prostratin induced a 41.7 ± 0.6-fold (mean ± SD) increased GFP signal relative to untreated cells, while 15 additional compounds induced at least 15-fold increases. These compounds included several frequently used control LRAs including (+)-JQ1, panobinostat, and vorinostat (Boehm et al., 2013; Rasmussen et al., 2013). They also included several that have been previously reported to reverse latency including AR-42, belinostat, bromosporine, CPI-203, dacinostat, I-BET-151, OTX015, PFI-1, and resminostat (**Table 1**) (Mates et al., 2015; Rasmussen et al., 2013; Pan et al., 2017; Liang et al., 2019; Shan et al., 2014; Li et al., 2019; Lu et al., 2016; Lu et al., 2017; Palmero et al., 2019). However, we also identified three previously unreported LRAs including the HDAC inhibitor CUDC-101 (Cai et al., 2010), which at 10 µM induced an 18.7 ± 0.8-fold increased fluorescence, the BET bromodomain inhibitor molibresib (I-BET-762; 29.9 ± 2.5-fold) (Nicodeme et al., 2010), and the HDAC inhibitor quisinostat (JNJ-26481585; 50.3 ± 0.8-fold) (Deleu et al., 2019; **Table 1**). These three compounds also maintained latency reversal with dose-dependence in both microplate reader and flow cytometry-based methods (Figures 5B-C), indicating that the microplate reader-based method can successfully identify and validate dose-response profiles of novel LRAs.

**Figure 5.**
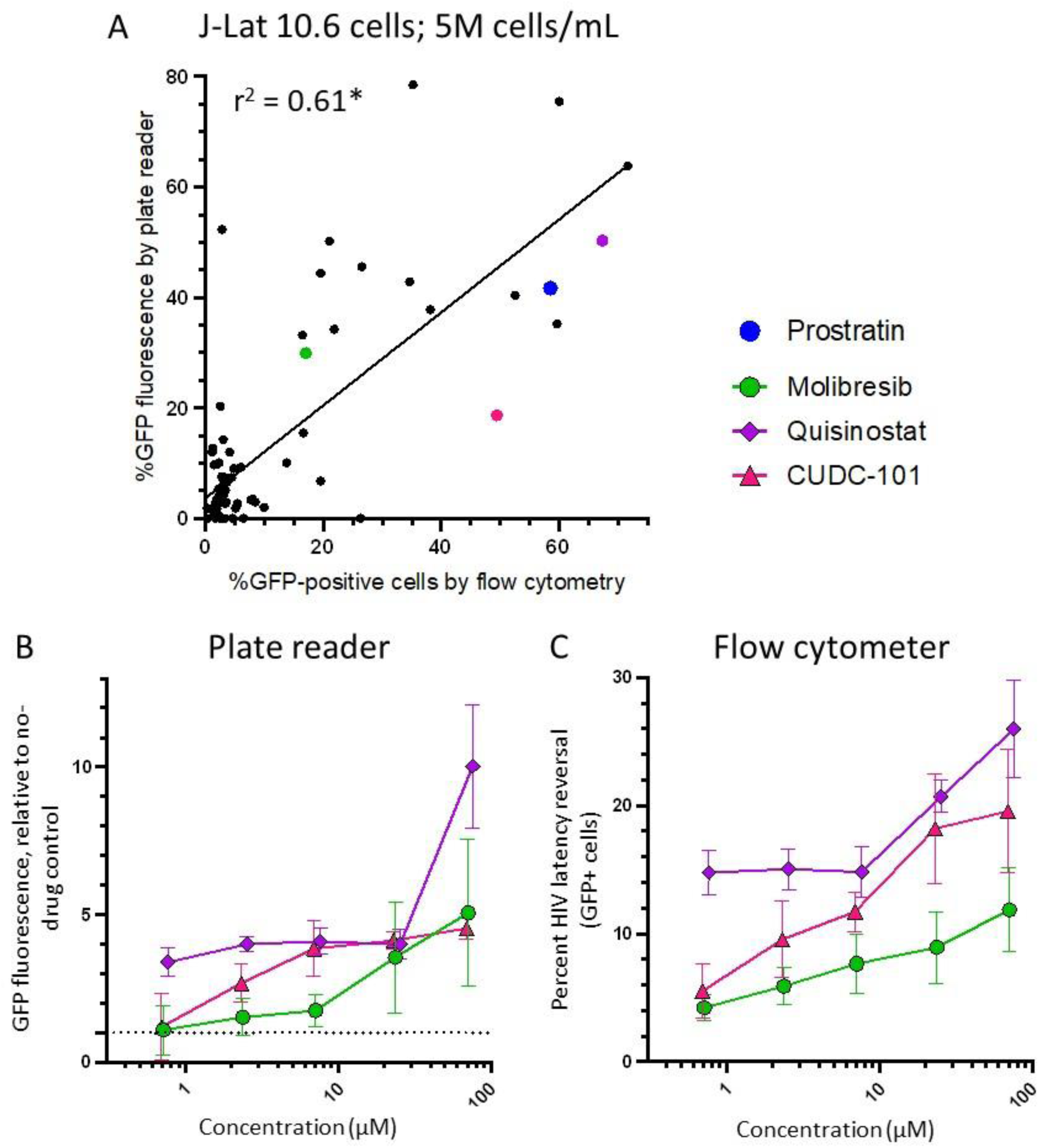
Discovery of novel LRAs from an epigenetic compound library. **A**, Correlations of microplate reading and flow cytometry data for 79 epigenetic modifiers screened at 10 µM. The blue dot denotes prostratin positive control, and other dots with colors denote 3 compounds that induce GFP expression at least 15-fold over untreated cells in the microplate reading-based method that have not been reported as LRAs. **B-C**, Dose response profiles of latency reversal due to molibresib, quisinostat, and CUDC-101 using the microplate reader-based method (**B**) and flow cytometry (**C**). In panel **A**, data are presented as the average of duplicates obtained by microplate reader method and flow cytometry. For panels **B-D**, data are presented as the mean ± s.e.m. from three independent experiments. *, linear regression p-value < 0.001.

**Table 1.**
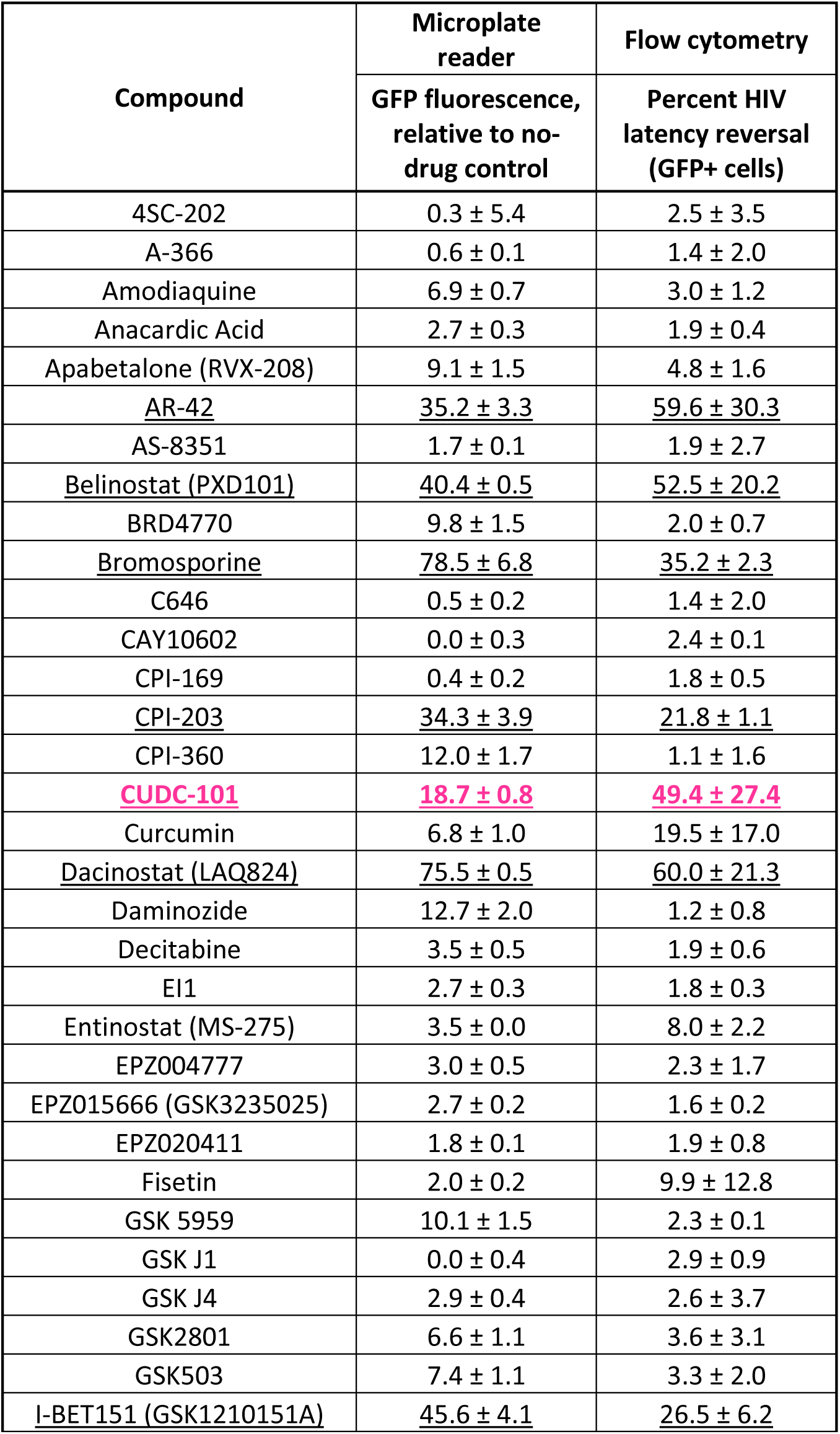
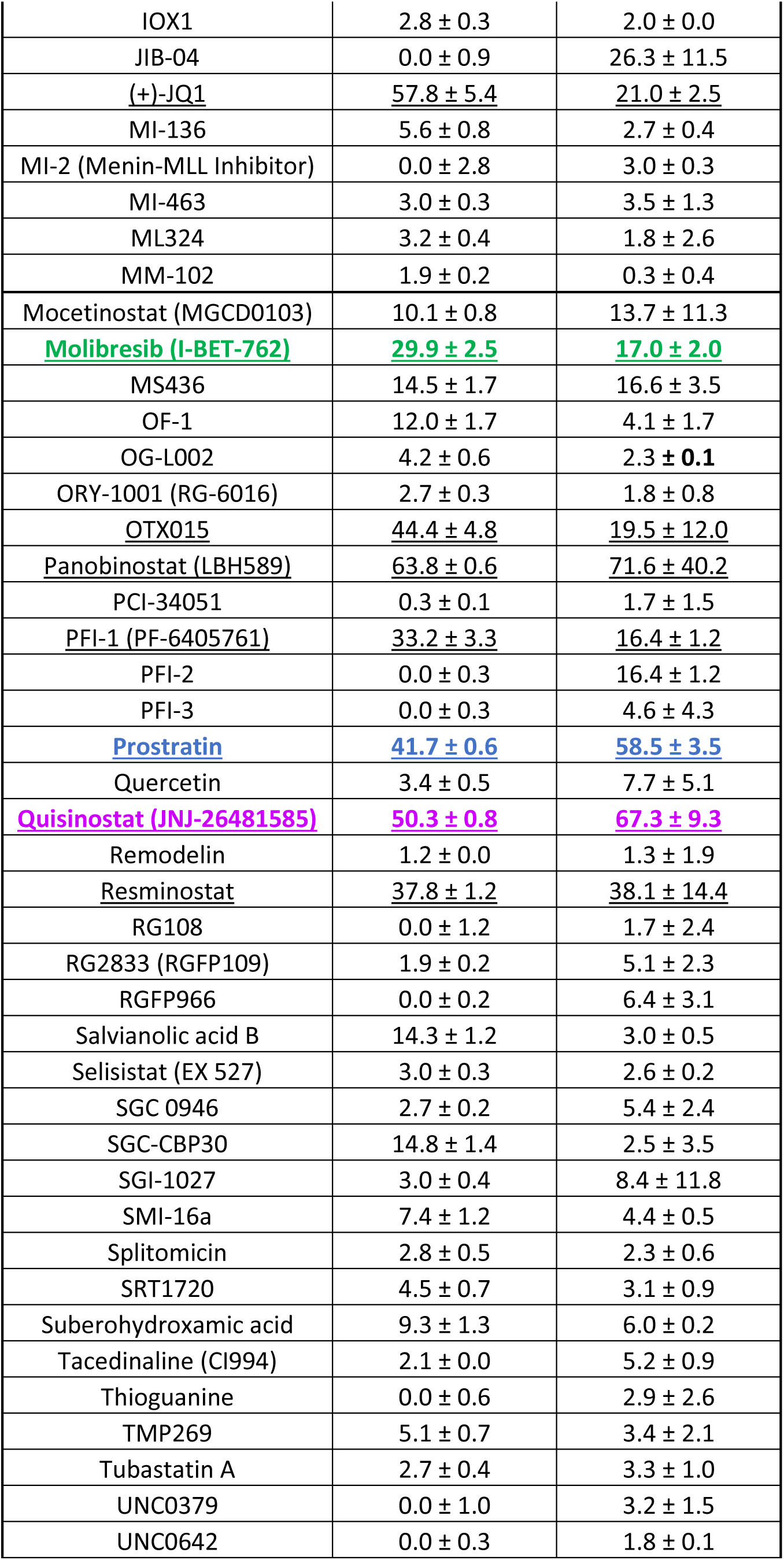
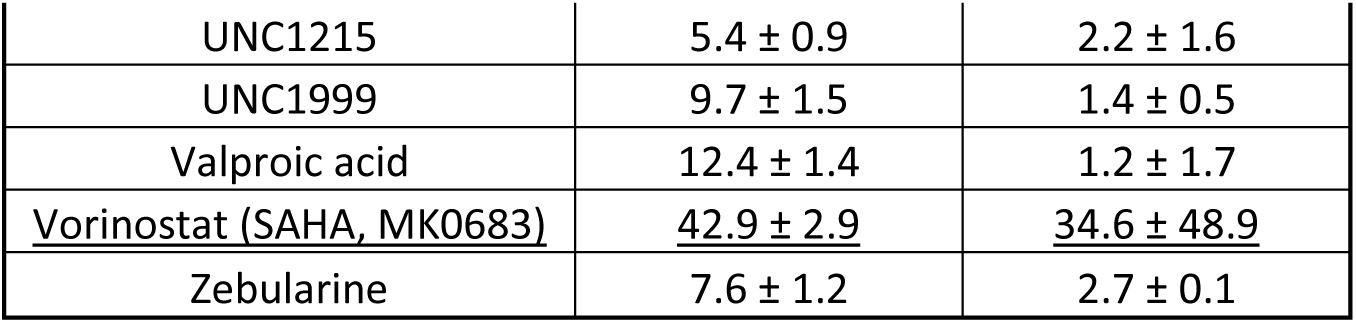
Latency reversal properties of epigenetic compounds assessed by microplate reader-based method and flow cytometry. LRAs that induce at least 15-fold GFP increase over untreated cells are underlined, while colors denote previously unreported LRAs.

## 4. Conclusion

Here we demonstrate the ability to identify and characterize novel LRAs in high-throughput format using J-Lat CD4+ T-cells and a standard microplate reader. The assay detects control LRAs with dose dependence, along with strong correlation and similar sensitivity to results obtained from flow cytometry, particularly using the highly responsive J-Lat 10.6 cell line but also to an extent in poorly responsive J-Lat 6.3 cells. We further demonstrate that this assay can be used to screen chemical libraries to identify novel LRAs, for example CUDC-101, molibresib, and quisinostat. Limitations of this assay include that compounds that induce cytotoxicity may present as false negatives at certain concentrations, as cytotoxicity will reduce overall GFP fluorescence which is otherwise circumvented in flow cytometry by assessing only the live gate subset of cells. Auto-fluorescent compounds would also be expected to appear as false positives, although these could be ruled out by subsequently incubating compounds with Jurkat cells which lack GFP fluorescence. Further, as J-Lat cells represent an *in vitro* model of HIV latency, new LRA hits will still require further validation such as measures of RNA induction or viral replication capacity after reactivation in primary CD4+ cells from study participants living with HIV. Nevertheless, this approach affords *in vitro*-based research on known and novel LRAs and mechanisms of HIV latency without the need for flow cytometry, thereby providing additional access to discovery-stage HIV cure research for highly resource-constrained laboratories in the absence of flow cytometry infrastructure and expertise.

## 5. Abbreviations

ART: antiretroviral therapy
CC_50_: half-maximal cytotoxic concentration
HDAC: histone deacetylase
LRA: latency-reversing agent
MTT: 3-(4,5-dimethylthiazol-2-yl)-2,5-diphenyltetrazolium bromide tetrazolium.

## 6. Author contributions

C.E.N.: Investigation, laboratory analysis, drafting of the original manuscript. U.S.: Investigation, laboratory analysis, drafting of the original manuscript. K.L.: Investigation, laboratory analysis. J.C.: Investigation, laboratory analysis. J.M.S.: Supervision, review & editing. L.J.M.: Funding acquisition, review & editing. R.N.N.: Supervision, review & editing. S.N.E.: Supervision. F.N.K.: Funding acquisition, supervision, writing, review & editing. I.T.: Funding acquisition, conceptualization, supervision, writing, review & editing.

